# Neighbour Typing Using Long Read Sequencing Provides Rapid Prediction of Sequence Type and Antimicrobial Susceptibility of *Klebsiella pneumoniae*

**DOI:** 10.1101/2025.09.03.673989

**Authors:** Mabel Budia-Silva, Amanda C. Carroll, Hiren Ghosh, Allison McGeer, Tommaso Giani, Gian Maria Rossolini, Karel Brinda, William P Hanage, Hajo Grundmann, Derek R MacFadden, Sandra Reuter

**Affiliations:** Institute for Infection Prevention and Control, Medical Centre – University of Freiburg, Freiburg, Germany; Faculty of Biology, University of Freiburg, Freiburg, Germany; The Ottawa Hospital Research Institute, Ottawa, Ontario, Canada; Sinai Health System, Toronto, Ontario, Canada; Department of Experimental and Clinical Medicine, University of Florence, and Microbiology and Virology Unit, Florence Careggi University Hospital, Florence, Italy; Inria, Irisa, Univ. Rennes, Rennes, France; Center for Communicable Disease Dynamics, Harvard T. H. Chan School of Public Health, Boston, Massachusetts, United States

**Keywords:** RASE, prediction, antibiotic, susceptibility, *Klebsiella pneumoniae*

## Abstract

The rapid genome-based diagnostic approach of long read sequencing coupled with neighbor typing offers the potential to improve empiric treatment of infection. However, this approach is still in development, and clinical validation is needed to support its use. In this study, we present an assessment of a neighbour typing method (RASE - resistance associated sequence elements) to predict lineage or sequence type (ST) and antimicrobial susceptibility in real time for *Klebsiella pneumoniae* sensu lato. We analysed the initial reads generated during the early phase of long read sequencing from pure culture (n=99), mock communities (n=20) and metagenomic samples (n=20). RASE accurately identified 69.7% and 70% of STs in pure culture and metagenomes, respectively, and identified the STs of the isolates representing the highest proportion in mock communities. Regarding antimicrobial susceptibility prediction, the probability of susceptibility increased to 72% (95% CI 63%-80%) across all tested antibiotics, when RASE predicted susceptibility, and decreased probability of susceptibility to 8.9% (95% CI 6.4%-9.6%) when was indicative of a resistant phenotype. Our study confirmed that genomic neighbor typing in *K. pneumoniae* sensu lato is capable of providing informative predictions of ST and antibiotic susceptibility in less than ten minutes (after the start of sequencing) with 200-500 reads.

**IMPORTANCE:** The growing burden of antimicrobial resistance is leading to high rates of mortality and morbidity worldwide. This situation has made the selection of empirical antibiotic therapy challenging, due to the risk of treatment failure and the overuse of last-resort antibiotics. The development of new sequencing technologies is helping to reduce the waiting time for a microbiological diagnosis, providing information in the early phase of bacterial infections, which could help improve clinical outcomes in a time of rising antimicrobial resistance. In this context, we assessed the performance of RASE (resistance associated sequence elements) in *Klebsiella pneumoniae*, an opportunistic pathogen frequently associated with nosocomial infections, which can rapidly acquire antibiotic resistance genes. Thus, in our study we provide insights that may aid in the validation of RASE for clinical use.

## INTRODUCTION

The increasing rate of antimicrobial resistance (AMR) is leading to a dearth of therapeutic options for the treatment of bacterial infections (1,2). In 2019, an estimated 1.27 million deaths worldwide were attributed to bacterial AMR, with *Klebsiella pneumoniae* ranked as the third leading pathogen associated with resistance-related mortality (3). Among these cases, *K. pneumoniae* producing extended-spectrum beta-lactamases (ESBL) and/or carbapenemases (CPE) were responsible for an estimated 50,000 and 100,000 deaths (3). Many of these multidrug-resistant *K. pneumoniae* are distributed across diverse clonal lineages (4), however, only some specific clones such as sequence type (ST) 258/512, ST11/347/420, ST101, ST307, ST15 and ST147 are disseminated globally, making a major contribution to AMR dissemination (5).

To address the expansion of antibiotic-resistant organisms, the World Health Organization has developed a global action plan (6). Many measures have been explored, including the improvement of diagnostic tools to reduce the time required to determine the etiological agent of an infection (6). Given the extended waiting times for detailed pathogen identification and the urgent need for antibiotic therapy in cases of systemic infections (sepsis, meningitis, etc), broad-spectrum antibiotics are used as empirical treatment in order to cover more potential pathogens (7). A rapid microbiological diagnosis could improve antimicrobial stewardship by an earlier selection of accurate treatment, decreasing the incidence of morbidity and mortality associated with bacterial infections (8), and reducing unnecessary antibiotic selective pressure arising from the use of broad spectrum agents (9).

Currently, the average time to obtain a microbiologic diagnosis using the classic culture-dependent tools is around ∼54 hours (10). New sequencing techniques such as long read sequencing (11), together with genomic neighbour typing, can reduce this waiting time to 4 hours from the moment the clinical sample is collected (12). Genomic neighbour typing allows for rapid prediction of lineage and antimicrobial susceptibility of pathogens from clinical specimens by comparison with known and characterized isolates. This method is a two-step algorithm, which first matches sequences of samples generated in real time against a database of reference genomes with a known sequence type, reference phylogeny and associated antimicrobial susceptibility phenotype, and thereafter predicts the probable phenotype based on the closest match and the matching quality. Since closely related isolates typically share similar properties, this provides a reliable heuristic for rapidly inferring the phenotype of the query pathogen. To enable this approach, the software application resistance-associated sequence element (RASE) was created to compare the *k-mer* content of nanopore reads to reference genomes, then calculate similarity weights (13). It predicts the phenotypic profile by identifying the best-matching lineage and comparing it with resistant and susceptible neighbors to determine a susceptibility score (12). RASE was evaluated previously on *Streptococcus pneumoniae* and *Neisseria gonorrhoeae* (*12*), demonstrating the ability to predict their susceptibility profile within ten minutes of starting sequencing. In this study, we assessed RASE on *K. pneumoniae* because of its critical importance to public health. We first developed a RASE database and then evaluated its accuracy, sensitivity and specificity in lineage calling, and explored different approaches to identify strains and resistance genes during sequencing.

## RESULTS

### Building a K. pneumoniae sensu lato reference Database for RASE

A reference database comprising genome sequences, sequence type (ST) and antimicrobial susceptibility data for 8 antimicrobial agents of 1511 *K. pneumoniae* sensu lato strains from the EuSCAPE study (14) was built using RASE software. The selection was based on isolates that were recovered from several European countries, thereby covering an extensive geographical range and ensuring representation of a large and diverse population. The database comprises strains belonging to 319 different STs within 14 clonal lineages, also including *Klebsiella variicola* and *Klebsiella quasipneumoniae*. The predominant STs were high-risk clones ST512, ST11, ST101, ST15, with more than 100 isolates aside from ST258 with 72 isolates, and covered strains with variable antimicrobial susceptibility profiles, ranging from susceptibility to all tested antibiotics to complex multidrug resistant profiles (Fig 1, Supplementary table 1).

**Figure 1.**
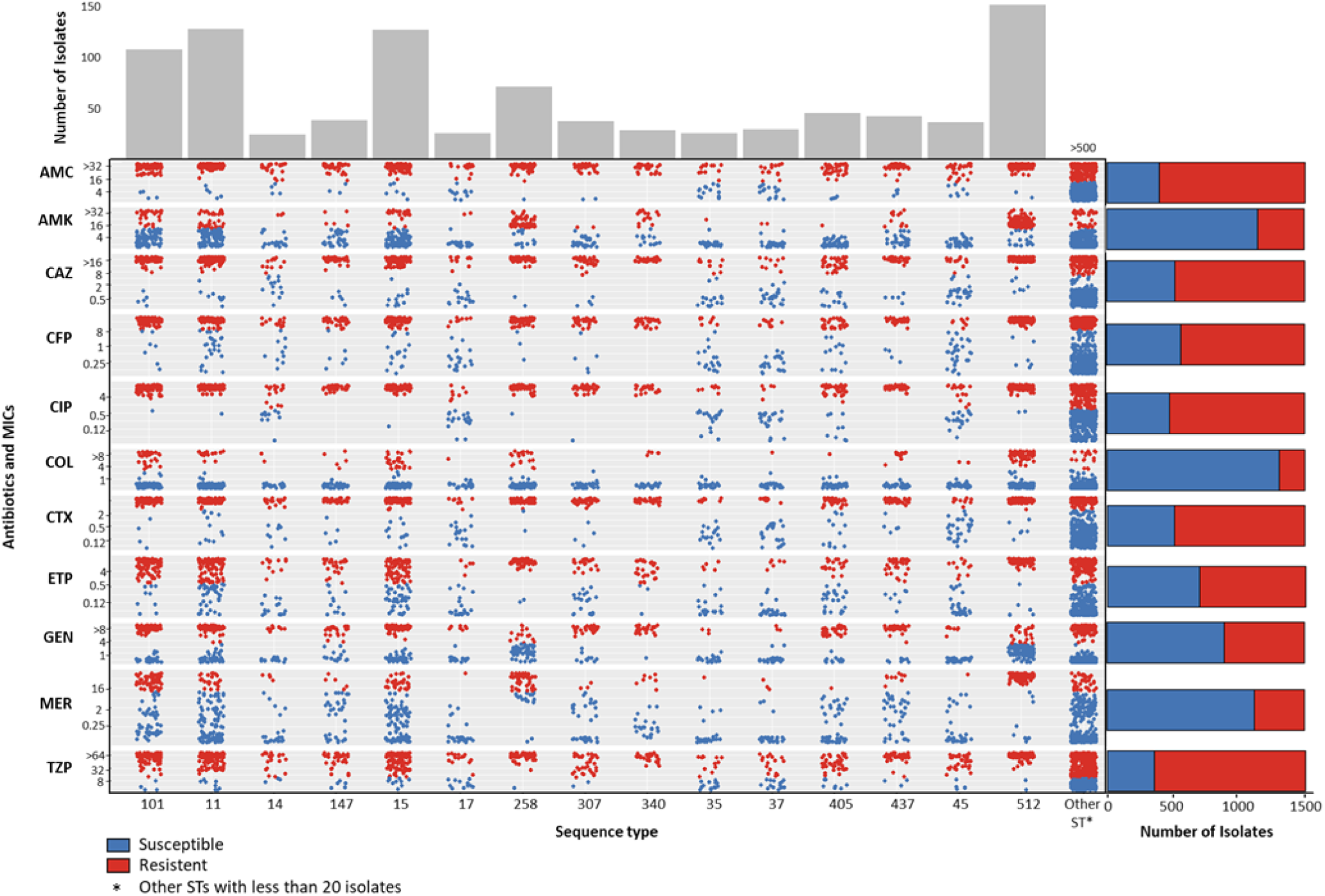
Overview of *Klebsiella pneumoniae* sensu lato RASE database, the dots represent an isolate coloured by their antimicrobial susceptibility for antibiotics in x, and are grouped in y according to the Sequence Type. Barplosts show the number of isolates susceptible, and resistant by antibiotic.

Strains belonging to the high-risk clones (15) were typically resistant to most antibiotics when compared to other clones (Fig.1), with more than 80% resistance to ciprofloxacin and piperacillin-tazobactam. Similarly, a high percentage demonstrated resistance to beta-lactams, with the exception of ertapenem, where susceptibility rates reached 50% and 56% in ST307 and ST405, respectively. For meropenem, resistance rates exceeded 94% in ST512, 75% in ST258, and 56% in ST101. Regarding aminoglycosides, more than 80% of ST258 and ST512 were resistant to amikacin, while a similar percentage of the same clones were susceptible to gentamicin. Notably, over 60% of all high-risk clones were susceptible to colistin. Other STs did not demonstrate a consistent susceptibility pattern.

### Comparison of the Performance of Read Extraction Methods and Evaluation of Preprocessing Tools

As the ultimate goal is to predict susceptibility/resistance from a small number of first reads obtained, we evaluated two strategies for read extraction (nanotimes, first fastq) and one strategy preprocessing (nanotimes+porechop+filtlong [NPF]) for improved quality, as well as their impact on the resulting prediction time. We extracted the first reads generated at the beginning of nanopore sequencing from 88 pure cultures (see Methods). We compared the results obtained by RASE, specifically the phenotypic prediction of eight antibiotics, with the MIC values (considered as true values), to calculate performance metrics, including sensitivity and specificity of susceptibility for each antibiotic. These values (Fig 2a) indicate a comparable overall performance. The sensitivity and specificity values were similar across approaches, although the NPF approach demonstrated the highest median sensitivity (0.75) and specificity (0.65), confirming improved data quality after applying Porechop and Filtlong. No statistically significant differences were found (Friedman test, sensitivity: χ² = 4.36, p = 0.113; specificity: χ² = 1.93, p = 0.381). Post-hoc Wilcoxon tests also showed no significant pairwise differences after multiple testing corrections (p >0.05 for all comparisons; Supplementary Table 2).

**Figure 2.**
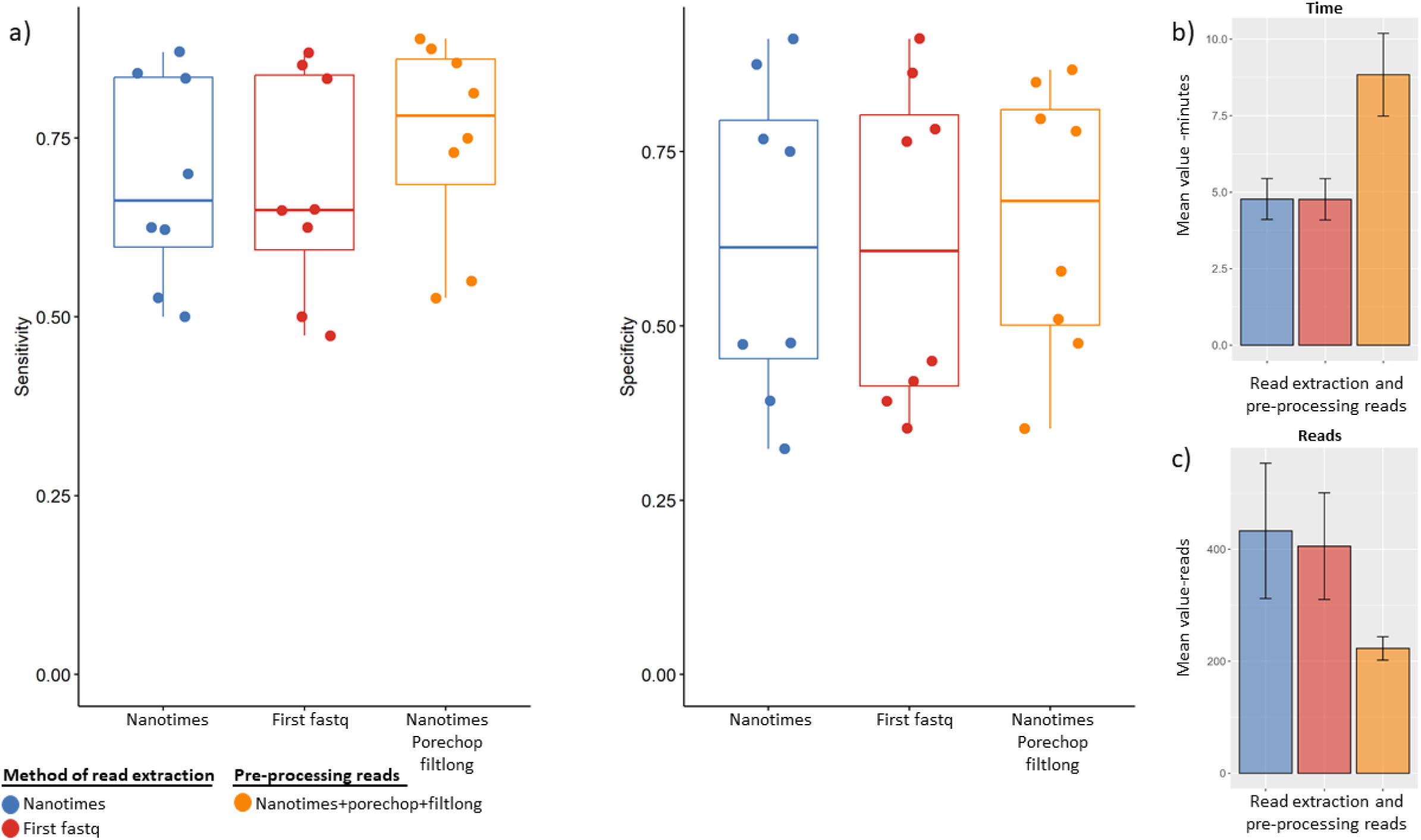
Comparative performance of two read extraction strategies and pre-processing reads in terms of phenotypic prediction accuracy, time to stable call, and number of reads required. a) Sensitivity (left) and specificity (right) of predictions for eight antibiotics were evaluated across two read extraction strategies: nanotimes (blue), first fastq (red), and nanotimes+Porechop+Filtlong (NPF, orange). b) Time (in minutes) and c) number of reads to achieve a stable call.

In addition, we analyzed computational performance regarding the time and number of reads required to achieve a stable call. As shown in Fig 2b, nanotimes and first fastq required approximately 5 minutes on average (mean = 4.77 minutes, 95% CI: 4–5.44 minutes), while the NPF approach required considerably more time (mean = 8.84 minutes; ∼10 minutes when considering the upper range). Regarding the number of reads needed (Fig 2c), nanotimes achieved the stable call using the highest number of reads (mean = 433), followed by first fastq (mean = 405), and *NPF* (mean = 223).

Taken together, these findings highlight that while all three approaches yield similar sensitivity and specificity for phenotypic prediction, the nanotimes approach demonstrates slightly better sensitivity, albeit requiring more reads. Based on a favorable balance between prediction accuracy and processing time, nanotimes was selected as the default method for subsequent analyses.

### Performance for Lineage Calling for Pure Cultures, Mock Metagenomics and Metagenomic Samples

To validate RASE lineage calling, we evaluated its performance across three groups of samples: pure cultures (n=99), mock communities (n=20) with known composition, allowing assessment under mixed-population conditions, and metagenomic samples (n=20). For pure culture isolates (Fig. 3a), RASE correctly predicted the ST in 69.7% of cases, with 82.6% of those recognized as the best match. In 14.1% of cases, the wrong ST was predicted, however 35.7% of these were in the same clonal lineage and therefore closely related (Fig 3a). For 16.2% of isolates, the ST was not present in the database, thus RASE was unable to make a correct call, however, classification was possible at the level of the clonal lineage for 37.5% of those isolates. The concordance was calculated using Cohen’s Kappa test to assess the level of agreement between the true ST (MLST, short read data) and the ST predicted by RASE, the concordance was 0.823 excluding those isolates belonging to STs not present in the database, and when including the complete collection, the concordance was 0.686. Both had a *p* =0, suggesting that the observed agreement provides strong validation for the accuracy of RASE in predicting STs represented in the database and related ones.

**Figure 3.**
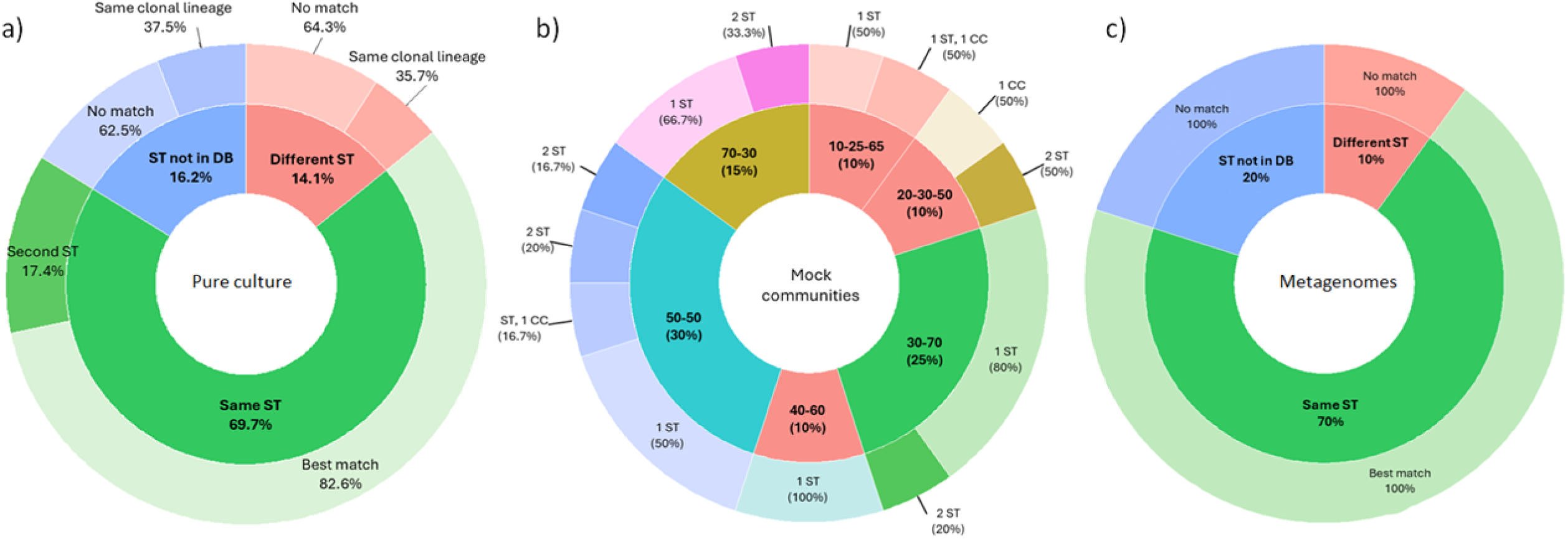
Validation of lineage calling of *K. pneumoniae* sensu lato RASE database across three conditions studied: a) pure culture b) mock communities and c) metagenomic samples.

Additionally, we verified the true genomic neighbor. For this analysis, we constructed three phylogenetic trees including isolates from our database and isolates used for validation (pure culture and isolates forming mock communities). Since ST258/512 was one of the most prevalent clonal lineage in our database, we reconstructed a separate tree focused on this lineage and we confirmed that the tested isolates were accurately predicted by RASE as belonging to the same ST (indicated by orange arrows in Supplementary Figure 1a). For those STs not present in the database, RASE correctly identified them as belonging to the same clonal lineage. Only in three cases did RASE predict a different ST. In the phylogenetic tree comprising the STs considered high-risk clones, RASE predicted the correct ST in most cases (Supplementary Figure 2a). The final tree, which includes the remaining STs, RASE also demonstrated accurate predictions. However, for several of these STs not represented in the reference database, RASE was still able to predict the ST of a closely related isolate (Supplementary Figure 3a).

To assess the effect of mixed samples, we prepared 20 mock communities: 15 with isolates of different STs and 5 with isolates of the same ST, yet distinct, i.e. not closely related, each subset with distinct ratios (Supplementary table 3). Among the subset with the same ST, the STs of 3 mock communities were accurately predicted, while the STs of the remaining mock communities were classified into the same clonal lineage. Within the mock communities with different STs, in the 50:50 ratio subset, one of the two STs present in the mock community was identified, while the second ST was identified as an alternative match. The other collections were mixed with different ratios, in which one of the STs was predominant. In these communities, RASE predicted the STs of the isolates with the highest proportion, except for two communities where the ST of the isolate with the highest proportion was recognized as a second match, and another case where the isolate with the lower proportion was identified as a second match (Table 1, Fig 3b).

**Table 1.** Summary of the accurately predicted lineage for mock communities.

|  | Lineage prediction | Ratio |  |  |  |  |  |
| --- | --- | --- | --- | --- | --- | --- | --- |
|  |  | 10-25-65<br>(n=2) | 20-30-50<br>(n=2) | 30-70<br>(n=5) | 40-60<br>(n=2) | 50-50<br>(n=6) | 70-30<br>(n=3) |
| 1 ST-high<br>proportion | Best match | 1 |  | 3 | 2 | 5 | 3 |
|  | Best match different ST<br>same CC | 1 | 1 |  |  | 2 |  |
|  | Second match | 1 | 1 | 1 |  | 1 |  |
| 2 ST-low<br>proportion | Best match | 1 | 1 | 1 | 1 |  |  |
|  | Best match different ST<br>same CC |  | 1 |  |  |  |  |
|  | Second match |  |  | 2 |  |  | 1 |
| 3 ST-lowest<br>proportion | Best match | 1 |  |  |  |  |  |
|  | Best match different ST<br>same CC | 1 |  |  |  |  |  |
|  | Second match |  |  |  |  |  |  |

Among the 20 metagenomic samples, 70% of STs were accurately predicted, regardless of the abundance of *K. pneumoniae*, identified as the best match. 10% were classified in another lineage and 20% were not present in the database (Fig 3c). The concordance of this evaluation was 0.62 and increased to 0.81 excluding those samples belonging to STs not present in our database. Based on the presented results, the best match and second match were often related, frequently belonging to the same clonal lineages; this outcome was consistently influenced by the representativeness of the database.

### Phenotypic Prediction from Pure Cultures and Mock Metagenomics

The RASE performance to predict antibiotic susceptibility phenotype was assessed using the same datasets for the evaluation of lineage prediction, however, samples lacking available MIC data were excluded, resulting in a total of 88 pure culture isolates and 10 mock communities that were analyzed. The overall sensitivity and specificity for susceptibility across eight antibiotics in pure culture were 0.71 (95% CI 0.67-0.75) and 0.68 (95% CI 0.65-0.73), respectively. Analyzing only the isolates with a lineage score >0.5 (n=25), both parameters increased to 0.91 (95% CI 0.84-0.97) for sensitivity and 0.72 (95% CI 0.64-0.80) for specificity. Certain antibiotics, such as amikacin and meropenem, exhibited a sensitivity of 1, while ciprofloxacin demonstrated a specificity of 1 (Supplementary Figure 4).

After RASE predicted a phenotype in pure cultures as susceptible or resistant, we measured the probability of susceptibility to individual or all antibiotics. This was compared to the empiric treatment thresholds of 80% for mild infections and 90% for moderate to severe infections (16). Across all antibiotics, the baseline probability of susceptibility was 44%, but it increased to 72% (95% CI 63%-80%), when RASE predicted a susceptible phenotype and decreased to 8.9% (95% CI 6.4%-9.6%) when RASE predicted resistant phenotype. With a probable susceptible call from RASE, each tested antibiotic exceeds the pre-test probability value. Nevertheless, ciprofloxacin improved to a 100% along with cefotaxime and colistin, which have post-RASE susceptibility probabilities exceeding 80% (Fig 4), suggesting that RASE predictions for susceptibility are more likely to be accurate.

**Figure 4.**
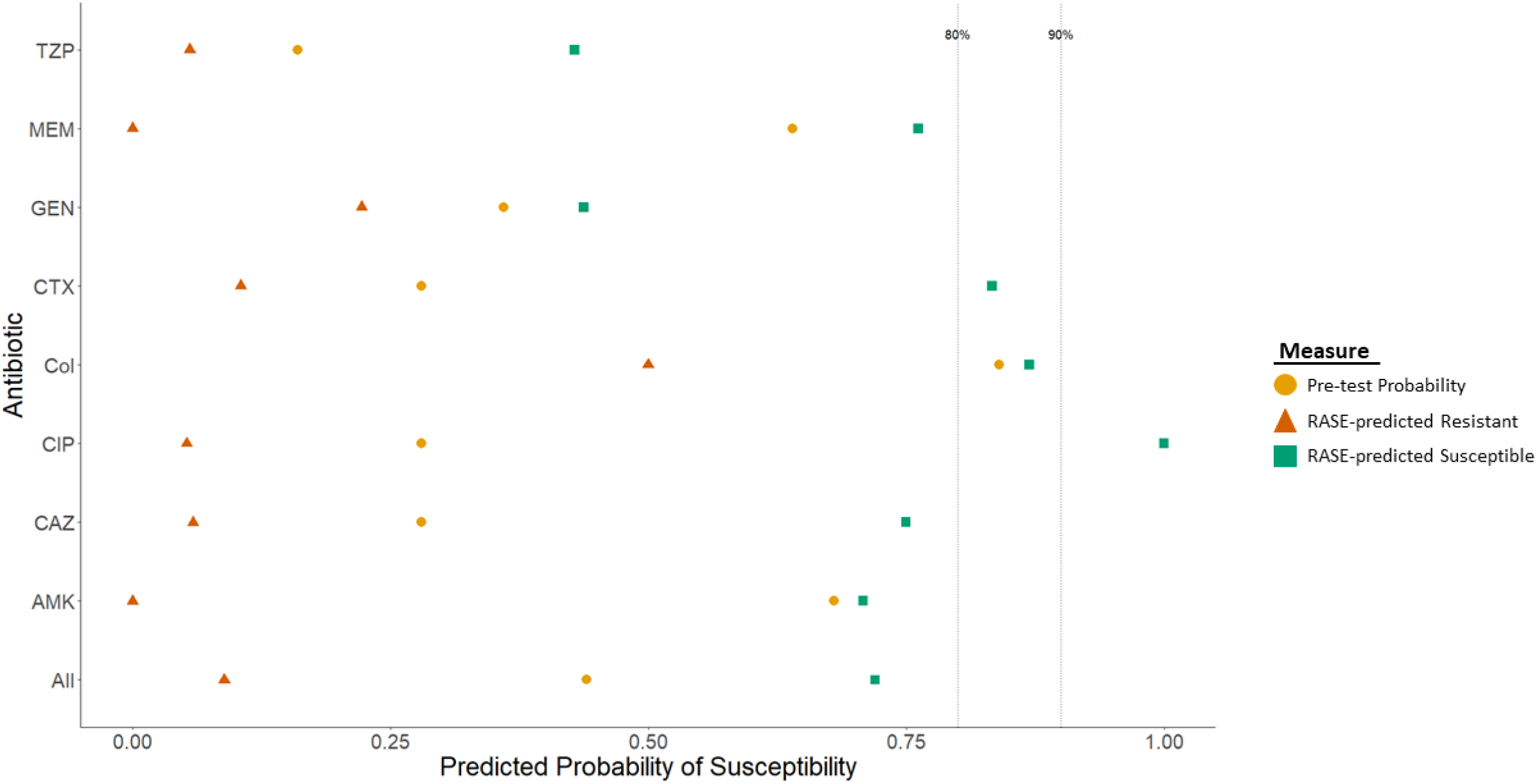
Probability of susceptibility of eight antibiotics individually and together predicted by RASE in 25 pure culture samples (Ls >0.5), shapes and colors differentiate the pre-test probability in yellow, predicted non-susceptible in orange and green the predicted susceptibility.

In this step, we evaluated RASE performance to predict antibiotic susceptibility phenotype in 10 mock communities as well, which were prepared with ratios of 50:50 (n=4), 30:70 (n=3) and 70:30 (n=3). The selection of isolates was based on their antibiotic susceptibility phenotype, one of the isolates of each mock community was multidrug resistant (R) while the other was multidrug susceptible (S) to eight antibiotics including amikacin (AMK), gentamicin (GEN), piperacillin-tazobactam (TZP), cefotaxime (CTX), ceftazidime (CAZ), meropenem (MER), ciprofloxacin (CIP) and colistin (COL). The mock communities 12 and 17 were prepared using the same isolates. Community 12 was composed at a 50:50 ratio, while 17 used a 30:70 ratio, the resistant isolate being present in higher proportion. In both cases, the best match was identified with the same sample. For communities 13 and 20, we similarly mixed two of the same isolates in a 50:50 or 70:30 ratio respectively, the best match corresponded to a susceptible isolate for these two collections. For communities 15, 16, and 19, RASE accurately predicted the susceptibility of the susceptible isolate at the 50:50 ratio. When the resistant isolate was present in a higher proportion, RASE identified the best match as a multidrug-resistant isolate. Conversely, when the susceptible isolate was in higher proportion, RASE identified as best match an isolate only resistant to one antibiotic. In most cases of our mock collections, RASE tended to predict the isolates present in the highest proportion (Table 2).

**Table 2.**
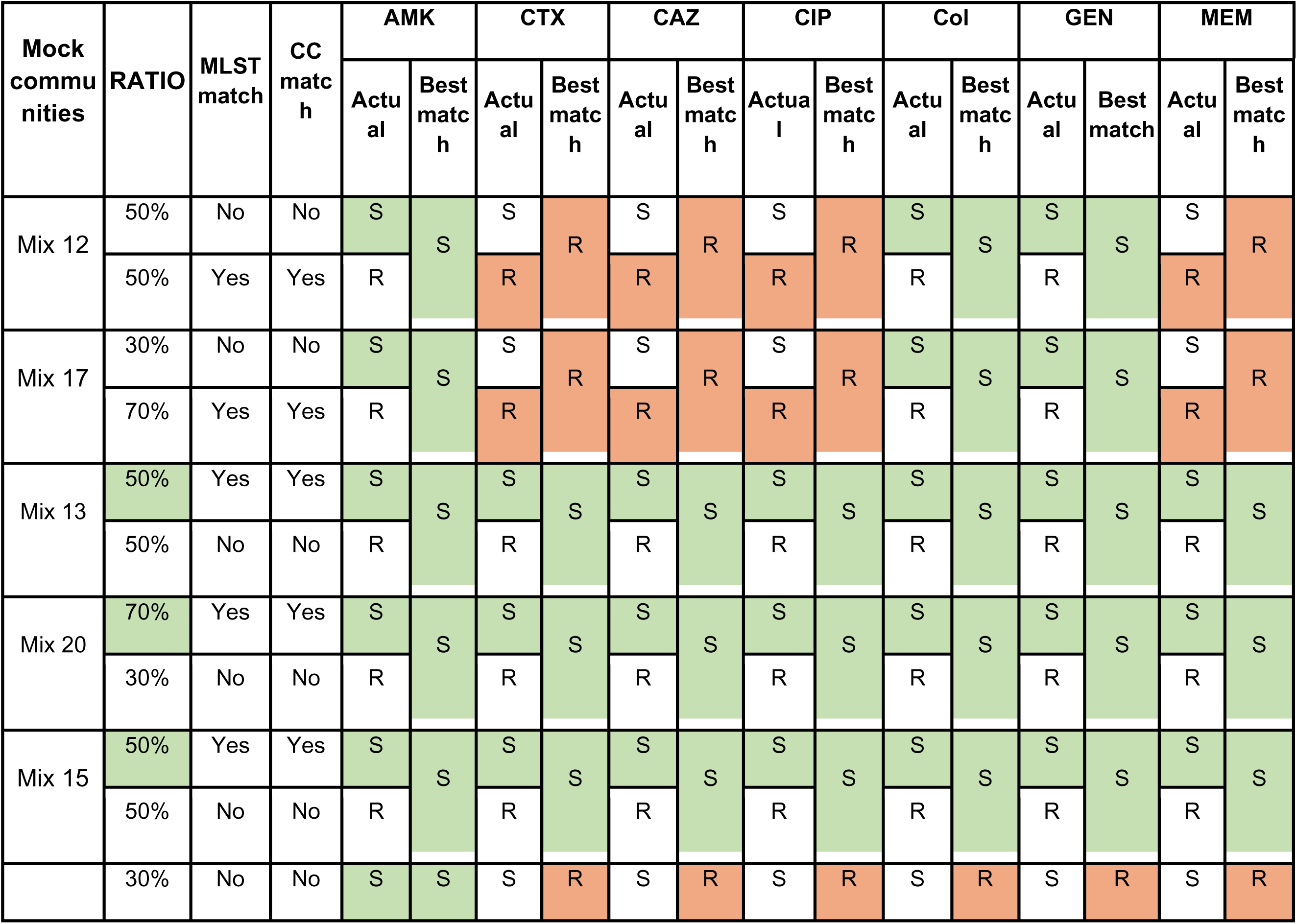
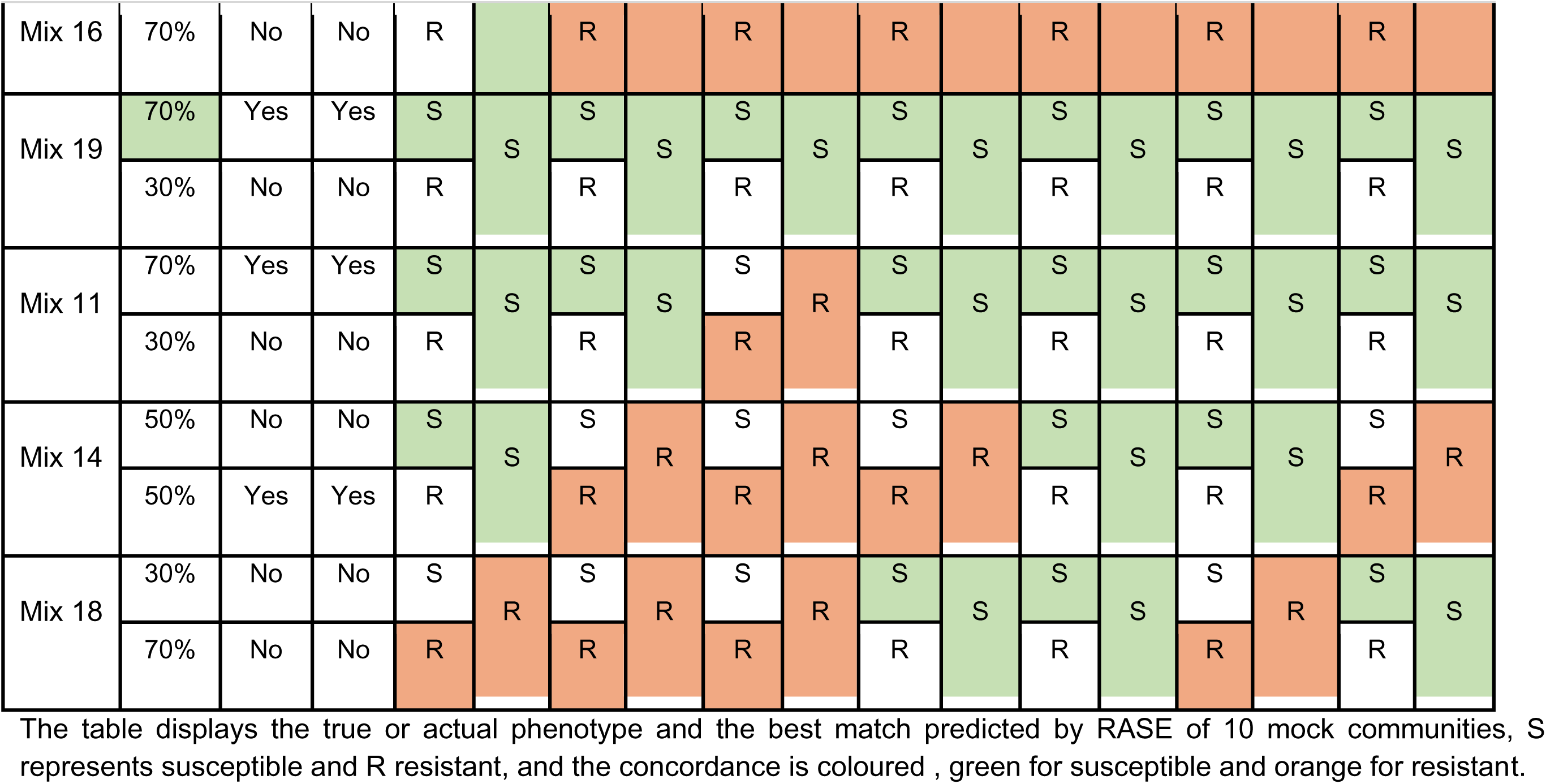
Predicted phenotype for mock communities. The table displays the true or actual phenotype and the best match predicted by RASE of 10 mock collections, S represent susceptible and R non-susceptible, and the concordance is coloured green for susceptible and orange for non-susceptible.

After we verified the true neighbors in lineage prediction, we contextualized RASE results for phenotypic prediction with the phenotypic profiles of database isolates, across the three phylogenetic trees. The pure culture isolates and isolates forming mock communities were grouped according to their ST, and the predicted antibiotic phenotype were similar to antibiotic phenotype of the isolates in the same cluster. However, isolates belonging to high-risk clones had more consistent patterns of resistance (Supplementary Figure 1c, 2c, 3b).

### Comparative Analysis of RASE performance with Gene-based predictive Methods

In order to compare RASE performance with gene-based predictive methods, we employed datasets from previous studies that analyzed AMR genes after 15 hours of sequencing. These samples were recovered in the USA (n=40) (17) and Germany (n=2) (18). The reported metrics show high sensitivity with the gene-based methods, however, the reported specificity was low for all or individual antibiotics. RASE demonstrated values over 0.9 for sensitivity and in most cases the specificity was higher for RASE (Supplementary Figure 5a) than for the gene-based method. Additionally, we evaluated the probability of susceptibility for 5 antibiotics, for all of them the values increased when RASE predicted susceptibility (Supplementary Figure 5b).

### Species Identification and Susceptibility Prediction in Metagenomic Samples

As the next step in our analysis, we aimed to assess the applicability of RASE in the metagenome of neonates. For this purpose, we retrieved 20 metagenomic samples collected within the framework of a study “Tracking the Acquisition of Pathogens In Real time” (TAPIR) which recovered nasal and anal swabs from premature neonates in the intensive care unit. We extracted the reads as described above, analyzing only the initial reads generated at the beginning of the sequencing process, and assessed whether this amount was sufficient to identify bacterial species. For this purpose, we employed Kraken2 in conjunction with Bracken to confirm the presence of *K. pneumoniae* and the bacterial proportions within 20 metagenomes. The results revealed consistent proportions of bacteria compared to those identified through the complete 72-hour sequencing analysis (Fig 5, Supplementary table 4).

**Figure 5.**
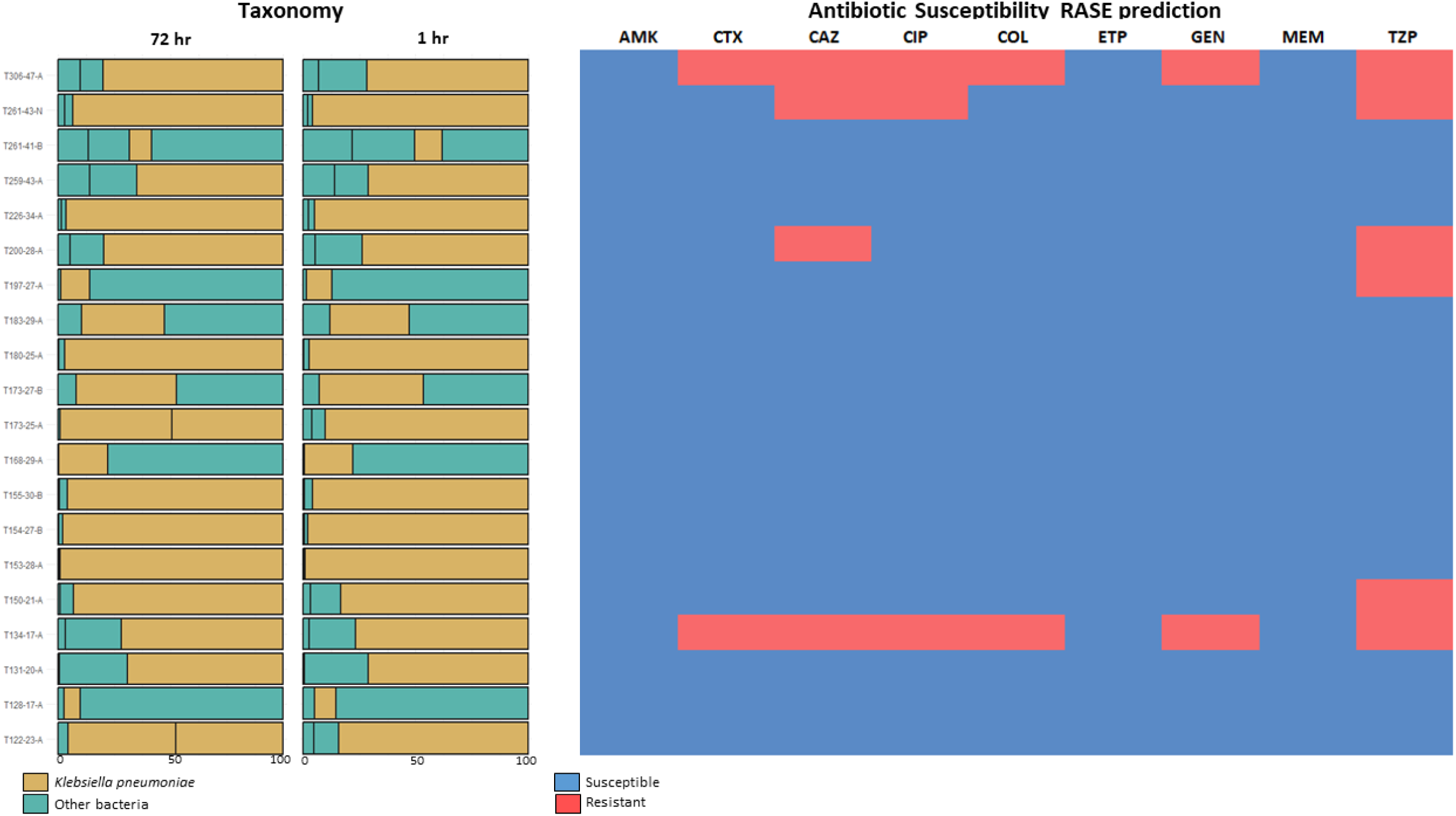
Analysis of metagenomic samples comparing the abundance after 72 hours and 1 hour of sequencing is shown in the first and second bartplots. The abundance of *K. pneumoniae* (Bracken analyses) is coloured in mustard yellow and in green other bacterias, considering that only the three most abundant bacteria per sample are displayed. Antibiotic susceptibility is depicted in the following columns, blue represents susceptible and red resistance isolates.

As expected for this sample cohort, the predicted susceptibility by RASE indicates that most of the samples (n=14) are susceptible to all tested antibiotics, while two samples are only susceptible to MER, AMK and Col (Fig 5).

## Discussion

The timeframe required to obtain a microbiological diagnosis through culture-dependent techniques takes days (10), impacting the timely and appropriate treatment of severe and common bacterial infections (19). This delay not only hinders the immediate initiation of effective antibiotic therapy but also contributes to the rise of AMR. Whereas appropriate empirical treatment helps to control infections effectively, inappropriate ones exert unnecessary selective pressure on pathogens. We present a promising approach combining long read sequencing technologies with algorithmic neighbor typing (12), to guide empirical antibiotic decisions and significantly reduce diagnostic turnaround time. In this study, we demonstrated that this approach improves the prediction of antibiotic susceptibility in comparison with traditional diagnostic methods (Fig 4). Moreover, the inclusion of lineage or ST prediction provides additional epidemiological insights to inform clinical decision-making, notably in the event of an outbreak. Finally, we observed no statistically significant differences in predictive performance among the read extraction strategies evaluated, supporting the robustness of the proposed method regardless of the input preprocessing pipeline.

The *K. pneumoniae* sensu lato used in this study are representative of a broader population at the continental level, as they include susceptible and resistant strains belonging to a wide range of STs, including globally circulating high-risk clones. This diversity enhances the relevance of the findings and supports the applicability of the database beyond regional contexts. In a clinical setting, a timely result would be highly beneficial. Thus, to ensure that the reads evaluated with RASE correspond to the initial reads during a sequencing run, we employed the nanotimes method for read extraction, which selects reads sequenced at the beginning of the sequencing process. Although the predictive performance of reads extracted by nanotimes and first fastq was comparable, nanotimes was preferred due to its alignment with the study’s focus on early diagnostic utility. Furthermore, our results demonstrate that the application of preprocessing tools such as Porechop and Filtlong (NPF) resulted in improved read quality but at the cost of increased processing time (Fig 2b), which may delay downstream analysis in time-sensitive contexts.

The validation of lineage calling and antibiotic susceptibility prediction using our database was conducted across three sample types: pure culture, mock communities and metagenomes. Concordance rates were high, with 0.82 in pure culture and 0.81 in metagenomic samples, underscoring the robustness of the approach. In mock communities containing more than one isolate, the pipeline consistently predicted the ST of the dominant strain (Table 1). In all tested conditions, RASE correctly identified the ST either as the best or second-best match. When the exact ST was not present in the database, RASE was still able to associate the isolate with its corresponding clonal lineage. In the absence of both ST and clonal lineage, the tool linked the strain to its nearest phylogenetic neighbor (Supplementary Fig 1-3). These results show the importance of maintaining a representative and frequently updated database, particularly at the regional level, to ensure accuracy in lineage and phenotype prediction as novel clones continue to emerge. Neighbor typing can complement existing approaches to surveillance, but not fully replace them.

Regarding the prediction of antibiotic susceptibility phenotypes, our analysis yielded an overall sensitivity of 0.91 (95% CI 0.84-0.97) and specificity of 0.72 (95% CI 0.64-0.80) across eight antibiotics and an increase in the predicted probability of susceptibility was observed for all tested antibiotics, supporting their potential applicability in treatment decisions. Moreover, RASE was able to accurately predict the susceptibility profile of the dominant strain in mock communities, however, this was not consistent across all antibiotics tested (Table 2). These results support the utility of RASE, when paired with a suitable database, as a rapid tool for simultaneous lineage and phenotypic prediction using a limited number of reads—typically between 300 and 500—generated in under 10 minutes for *K*. *pneumoniae* (Fig 2b,c).

Our findings align with previous studies evaluating RASE for other bacterial pathogens, including *Streptococcus pneumoniae* and *Neisseria gonorrhoeae*, demonstrating prediction times under 10 minutes. In those studies, specificity reached 100% for both species, while sensitivity was reported at 91% for *S. pneumoniae* and 81% for *N. gonorrhoeae* (12). Additionally, applications of RASE to *Escherichia coli* and *Klebsiella spp*. in Canada have emphasized the necessity of using regionally curated databases to ensure predictive accuracy (20) .

The development and integration of novel technologies aim to enable rapid microbiological diagnostics. Among the strategies are methods based on the direct detection of genetic material or proteins, and these approaches have shown high sensitivity, but they still require some hours of sequencing, representing a limitation. In contrast, RASE, as demonstrated through our comparative analysis (Supplementary Fig 5-6) in this study, exhibited promising performance with significantly reduced times.

A known limitation of the RASE-based approach is its dependency on a species-specific reference database (12,20). This requirement may hinder performance when the causative pathogen is unknown at the time of sequencing, and thus an additional step to identify putative relevant pathogens in a sample is necessary prior to then selecting appropriate RASE databases. Another challenge arising from metagenomic sequencing from swabs is the microbial content of clinical samples. Some swab-based screening samples may contain low biomass, limiting the amount of microbial DNA available for analysis. In our study, pre-enrichment using BHI medium improved microbial yield but resulted in extended processing times. By contrast, biological samples such as blood, urine, and sputum can be sequenced directly without enrichment, as demonstrated in previous studies (21–23).

In summary, our study demonstrates that genomic neighbor typing using the *K. pneumoniae* RASE database enables rapid prediction of both sequence type or lineage complex (concordance: 0.82) and antibiotic susceptibility phenotype (sensitivity: 0.91, 95% CI: 0.84–0.97; specificity: 0.72, 95% CI: 0.64–0.80). This combined approach has the potential to significantly reduce the time to microbiological diagnosis, facilitating more accurate empirical antibiotic selection and contributing to improved treatment outcomes in *K. pneumoniae* infections.

## METHODS

### Reference Klebsiella pneumoniae sensu lato Database

The RASE reference database was generated using assemblies, genotypic multi-locus sequence typing and antibiotic susceptibility phenotype of *K. pneumoniae* isolates from the EuSCAPE study. This study aimed to describe the epidemiology of carbapenem resistant Enterobacterales, by recovering susceptible and resistant *K. pneumoniae* strains between 2013 to 2014 across Europe (n=1511 strains) (14). We included 1511 fasta files of good quality, considering their N50, length and number of contigs, and for which antimicrobial susceptibility testing had been defined (Supplementary table 1). Broth microdilution was performed by reference methodology (24) to determine minimum inhibitory concentration (MIC) (Supplementary table 1). We categorized the MIC values according to the European Committee on Antimicrobial Susceptibility Testing (EUCAST) v15 breakpoints (24), assigning to each strain its respective sequence type (ST) and an antibiotic-specific resistance category (susceptible or resistant) for the following antibiotics: AMK, GEN, TZP, CFP, CTX, CAZ, MER, CIP and COL.

### Datasets for Evaluation

Validation was performed with susceptible and resistant *K. pneumoniae* strains from clinical primary specimens and isolates. These included 99 pure cultures selected based on their resistance profiles and STs, as well as the availability of the data we needed to analyze, such as MIC values (isolates from EuSCAPE [n=51] (14), EURECA [n=40] (25) and a project containing isolates from Greece [n=8] (26)) (Supplementary table 5). 20 mock communities were prepared from selected pure cultures, following the criteria described below (Supplementary table 3). 20 metagenomic samples of anal and nasal swab recovered from neonates through the TAPIR project were chosen according to the presence and abundance of *K. pneumoniae* (Supplementary table 4). All samples were previously short read sequenced and the antibiotic susceptibility profiles are available for the majority, with the exception of metagenomic samples and 11 pure cultures. Therefore, all phenotypic prediction analyses were performed using only 88 pure cultures. The previously determined STs found using short read data and MICs values were updated based on EUCAST (24) v15, where “susceptible increased exposure” values were reinterpreted as susceptible.

### DNA Extraction, Library Preparation and Nanopore Sequencing

Pure culture isolates, including those used for the construction of mock communities, were cultured on blood agar and incubated overnight at 37°C, then species identification was performed using MALDI-TOF mass spectrometry, and a single colony was selected for subsequent DNA extraction. Metagenomic samples were enriched with BHI medium and incubated overnight at 37°C, 210 rpm in a proportion 1:2. To remove the human DNA, enriched samples were centrifuged at 10.000 rpm for 5 minutes, the microbial pellet was resuspended with 1 mL of clean dH_2_O at room temperature for 5 minutes and a final resuspension in 200 μL PBS 1X; 5 μL lysozyme and 5 μL lysoplastin.

DNA was extracted using the Roche High Pure Template Preparation Kit, and long-read sequencing was performed using the Oxford Nanopore Technology (ONT). Sequencing libraries were prepared using the ligation protocol SQK-LSK114 with native barcode kits (EXP-NBD114 and SQK-LSK 114) for R9 Chemistry, and SQK-NBD114-96 barcoding kit for R10 Chemistry in accordance with the manufacturer’s protocol including optional steps. DNA input and other measurements were calculated based on the assumption that 1 fmol is equivalent to 5 ng of DNA. Thus, for the final R10 library, 100 ng corresponded to 20 fmol. For the R10 protocol, the initial volume per sample was 11 μL without water dilution (for clinical samples). Optional NEBNext FFPE DNA Repair and DNA Control Sample were omitted during DNA repair and end-prep. In addition, Bovine Serum Albumin (BSA) was used to improve the sequencing performance as recommended in the protocol.

Final libraries were quantified using Qubit 4 Fluorometer, and loaded onto a FLO-MIN114 R10.4 flow cell type and sequenced on a GridION X5 Mk1 sequencing platform. Sequence data acquisition, real-time base-calling, and demultiplexing of barcodes were conducted using the graphical user interface MinKNOW (v23.11.7) and the dorado basecaller (v7.2.13).

Sequencing data has been deposited in the following ENA projects: Mock Communities PRJEB82665 (supplementary table 3), TAPIR metagenomic clinical samples subsampled to first 60min PRJEB96334 and pure cultures PREJB95992 (supplementary table 4). Long read ONT sequencing used to evaluate RASE and the reference database from projects Daikos Greece PRJEB58216, EuSCAPE and EURECA PRJEB96336 (supplementary table 5).

### Analysis of Culture Specimens

To evaluate long-read sequencing from pure culture, extraction of reads from isolates were initially performed based on the sequencing time (60 min) using nanotimes (https://github.com/angelovangel/nanotimes; first dataset). Along with that, the first fastq sequenced for each isolate was extracted (second dataset). Porechop (v0.2.4; https://github.com/rrwick/Porechop/tree/master) was used to trim off adapters and filtlong (v0.2.1; https://github.com/rrwick/Filtlong) to filter the long sequences with a minimum read length threshold of 1000, only for reads extracted with nanotimes (third dataset [NPF]).Therefore, three pure culture sets of sequences of each sample were analyzed with RASE (version 0.1.0.0).

### Analysis of Mock Metagenomics

A total of 20 mock communities were created to evaluate the ability of long read sequencing coupled with neighbour typing to predict ST and phenotype in the setting of controlled consortiums of bacterial strains. Isolates to be mixed were chosen based on their ST and antibiotic susceptibility phenotype. Each mock community corresponds to the DNA pool of two (n=16) or three isolates (n=4), with different proportions, resulting in six subsets 50:50 (n=6), 40:60 (n=2), 30:70 (n=5), 20:30:50 (n=2), 10:25:65 (n= 2) and 70:30 (n=2) (Supplementary table 3). At least one mock community of each subset was prepared with isolates of the same ST (n=5), except the subset 70:30. Only ten of these communities were used to evaluate the antibiotic susceptibility prediction, including the subsets 50:50 (n=4), 30:70 (n=3) and 70:30 (n=3). Each mock community was prepared with one multidrug resistant (R) and one multidrug susceptible (S) isolate to eight antibiotics AMK, GEN, TZP, CTX, CAZ, MER, CIP and COL. The 20 mock communities were sequenced using nanopore, after the first reads sequenced during the beginning of the sequencing of each mock community were extracted using nanotimes (60min). The number of reads differed between 2551 to 17205 (Supplementary table 3), and then these sequences were assessed with RASE.

### Analysis of Metagenomic Samples

Extracted reads employing nanotimes of metagenomic samples were mapped to the human genome (GRCh38) using Minimap2 (v2.24; using the map-ont parameter) (27) and human DNA contaminants were removed from reads using Samtools (v1.14) (28). Species identification was performed using Kraken2 (13). Microbial species abundance was estimated using Bracken (29). Subsequent analysis was done with RASE.

### Analysis with RASE

Rase algorithm is based on matching the nanopore reads to a reference database using Prophyle and increasing the weight of the most similar strains. First, the lineage is identified by finding the best match reference genome. The lineage score is calculated by comparing the two best-matching lineages. In the next step, the best match within the lineage is identified, and resistance is predicted from the nearest resistant and susceptible neighbors and the susceptible score is a result of the comparison of their weights. To use the software we follow the steps described in the documentation, then to analyze our results, we first define the stable call based on the fluctuation of the lineage score (LS), when it did not change by more than 0.2 for at least 100 reads, as LS was created as a potential indicator of confidence in lineage prediction. When we performed long read sequencing of isolates that were already contained within EuSCAPE (n=51) (14), we created a new database for each sample by removing the sample being evaluated.

### Analysis of external datasets

Additionally, we analyzed two external dataset from two previous studies (17,18) that assessed gene-based methods for identifying antimicrobial resistance (AMR) genes. We downloaded 42 genome assemblies from the ENA database and obtained the corresponding raw sequencing reads generated after 60 minutes of Nanopore sequencing using nanotimes. The reads were processed and analyzed with RASE to predict AMR profiles. We then compared the sensitivity and specificity of our predictions against those reported in the original studies to evaluate performance consistency.

### Statistical Methods and Visualization

The concordance between the true ST and the one predicted by RASE was assessed with the Cohen’ s Kappa test using the irr package (30) (v2.0.60). Friedman test was performed to compare the three read extraction methods. The performance of RASE predicting susceptibility and non-susceptibility was evaluated by calculating its sensitivity and specificity. Sensitivity was determined by measuring the proportion of true positive results. This was calculated as the number of true positives divided by the sum of true positives and false negatives. Specificity, conversely, was calculated by assessing the proportion of true negative results. It was calculated as the number of true negatives divided by the sum of true negatives and false positives. Plots were visualized using ggplot from Tidyverse (31) (v2.0.0).

### Phylogenetic analyses

Phylogenetic trees of the database and tested isolates were estimated using RAxML v8.2.12 (32) based on SNP alignments after mapping to reference genome MGH78578 (GenBank accession CP000647) and removal of recombinant regions using Gubbins v2.4.1 (33). Phylogenetic tree visualization was done with iToL (34).

### Ethical approval

This study has been approved by the local ethics committee (22-1039 and 22-1040).

## ACKNOWLEDGEMENTS

This project was funded by JPIAMR call (K-STaR 01KI1910), and the Federal Ministry of Research, Technology and Space (BMFTR), formerly the Federal Ministry of Education and Research (BMBF; TAPIR 01KI2018) funding to SR. KB was supported by the French National Research Agency (ANR) under Grant ANR-24-CE45-1226 (REALL project).We thank technical support by Leonardo Duarte dos Santos.

---

**Supplementary Figure 1** Phylogenetic tree of database isolates, a) pure culture and b) isolates forming mock communities within the ST258/512 clonal lineage, clades are coloured by ST. Arrows link each tested isolate to the match predicted by RASE. Orange arrows indicate predictions within the same ST, light blue for matches in a different ST within the same clonal complex and grey shows isolates of STs absent from the database but matched within the same clonal lineages. c) Phenotypic prediction by RASE of evaluation samples, along with the true antibiotic phenotype of database isolates for eight antibiotics.

**Supplementary Figure 2** Phylogenetic tree of database isolates, a) pure culture and b) isolates forming mock communities belonging to the high-risk clones. Each tested isolate is linked to its predicted match according to RASE results. Arrows colours: orange indicates matches within the same ST, light blue matches in a different ST within the same clonal lineage, and purple indicates matches with a different ST. c) Phenotypic prediction by RASE of evaluation samples, along with the true antibiotic phenotype of database isolates for eight antibiotics

**Supplementary Figure 3** Phylogenetic tree of remaining isolates belonging to different STs, the most prevalent STs are coloured. Orange arrows indicate matches within the same ST, light blue arrows denote matches in a different ST within the same clonal lineage, and grey arrows correspond to isolates of STs that are not present in the reference database but are matched with the closest isolate. c) Phenotypic prediction by RASE of evaluation samples, along with the true antibiotic phenotype of database isolates for eight antibiotics.

**Supplementary Figure 4** Phenotypic validation of the *K. pneumoniae* senu lato RASE database in 25 pure culture samples, stratified by sensitivity and specificity.

**Supplementary Figure 5 a)** Comparative analysis of RASE (blue) and gene-based methods (red) in terms of sensitivity and specificity across six different antibiotics. Each dot represents the predictive performance for a specific antibiotic. **b)** Probability of susceptibility of six antibiotics predicted by RASE in 42 external isolates, shapes and colors differentiate the pre-test probability in yellow, predicted resistant in orange and green the predicted susceptible.

**Supplementary Table 1** Reference database information, including quality control of assemblies, multilocus sequence types, and antibiotic susceptibility profiles for the evaluated antibiotics.

**Supplementary Table 2** Results of statistical test (Post-hoc Wilcoxon, Shapiro-Wilk and Friedman) used for comparing read extraction methods and pre-processing tools.

**Supplementary Table 3** Proportions of isolates forming mock communities and their true STs. Also includes RASE ST predictions and ENA accession numbers.

**Supplementary Table 4** Metagenomic samples data including true ST, RASE ST predictions along with bacterial proportions in culture, sequencing after 1hr and 72 hrs. ENA accession numbers are included as well.

**Supplementary Table 5** This table presents information of pure culture including true ST and antibiotic susceptibility profiles for evaluated antibiotics. Additionally provides ENA accession number and RASE predictions (stable call) of each sample.

